# CoSiDeX: A hyperspectral fluorescent protein resource for highly multiplexed imaging

**DOI:** 10.1101/2025.05.09.652935

**Authors:** Alejandra V. Patino, Henry Chapman, Joshua D. Ginzel, Joelle E. Sills, Herbert Kim Lyerly, Bruce W. Rogers, Joaquin E. Drut, Joshua C. Snyder

## Abstract

Fluorescent proteins (FPs) have revolutionized spatiotemporal observations in biology. Yet, the design of multiplexed assays remains constrained by limited spectral characterization and palette validation. Although over 1,000 FPs have been catalogued, systematic resources for characterizing their use in multiplexed approaches are lacking. Here we present a resource and methodology for selecting and decoding FPs in multiplexed imaging experiments. A library of forty-four FPs was built for rapid assembly into mammalian expression vectors and transposase-mediated integration. Hyperspectral imaging was performed for each FP and spectral space was characterized mathematically. To support experimental design and data interpretation, we developed the Cosine Similarity Decoder of XFP (*CoSiDeX*) toolbox, to predict spectrally resolvable FP palettes and decode and re-color hyperspectral images. Using this approach, we demonstrate live-cell imaging of 12 uniquely labeled clones. Our work offers a scalable platform for selecting optimal FP palettes for multiplex experiments, with broad utility across diverse biological systems and hyperspectral imaging techniques.

## Introduction

Biology is highly interconnected, dynamic, and spans various spatiotemporal scales. Yet, studying processes across these dimensions remains challenging. Single-cell technologies have addressed some of these challenges by enabling simultaneous inference of intracellular states and intercellular relationships^1–3^. However, these technologies are constrained by high costs, destructive workflows, and the inability to continuously monitor dynamic processes. For more than two decades, green fluorescent protein (GFP)^4^ and its derivatives have offered a way to study live dynamic biological processes across spatial and temporal dimensions^5^. Fluorescent protein (FP) based methodologies have facilitated critical discoveries in tumor biology^6,7^, immunology^8,9^, neuroscience^10^, and protein dynamics^11,12^. Yet, most studies continue to use single-reporter systems from a limited palette of FPs. As the number of biological processes studied in a single context increase, there is a concomitant need for an expanded repertoire of spectrally resolvable FPs that have been validated for combinatorial imaging.

While there are now more than 1000 unique FPs^13^ across the visible and near infrared spectrum (Figure 1a), the expansion of FP multiplexing has mostly relied upon advances in genetic engineering and light microscopy utilizing only a few spectrally resolvable FPs. These include combinatorial lineage tracing and unmixing approaches used in the Brainbow reporter system^14,15^ and its derivatives^16–20^, cellular compartment and protein specific FP expression^21^, and lineage tracing tools for studying cancer initiation and progression as in our previously published Cancer rainbow (Crainbow) model systems^22–25^. However, because a comprehensive analysis and standardized resource for validating spectral resolvability is lacking, the burden of identifying multiplexable FP palettes falls on the investigator. For instance, when Crainbow models were first developed, FP choice was based upon literature searches and best guesses were informed qualitatively by spectra, if available. Thus, FPs had to be cloned, expressed, and tested empirically^22^. Since then, resources like FPBase^13^ have provide a much-needed repository for collecting and maintaining a database for FP sequence, physical properties, and spectra. However, standardization of FP spectral resolvability remains a bottleneck in the development of FP-multiplexable technologies. Thus, generating models with non-optimal or incompatible FPs continues to be a major challenge.

**Figure 1.**
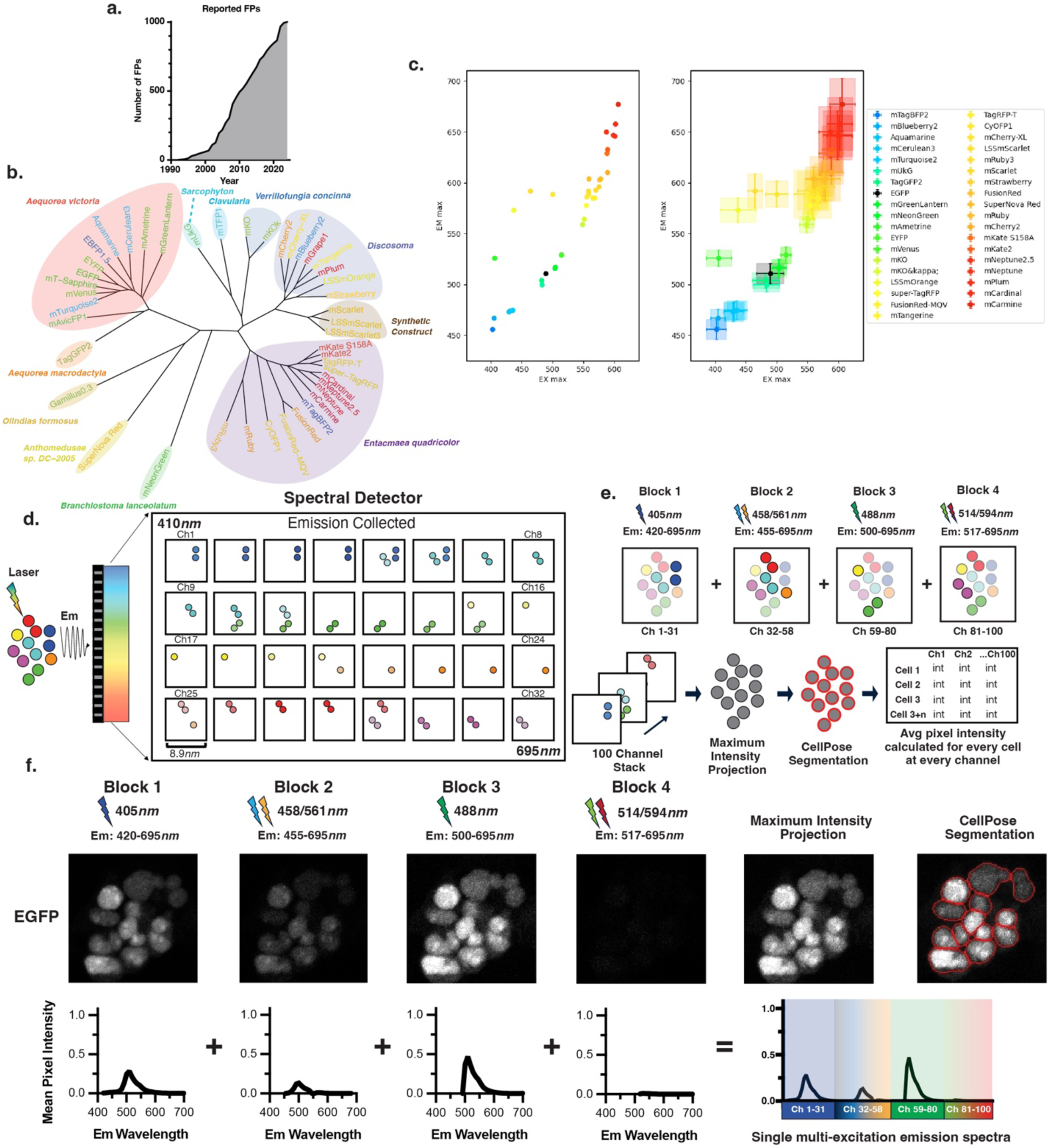
Fluorescent protein library and hyperspectral imaging approach. **a**. Growth in the number of catalogued FPs from 1994 to 2025. Data extracted from FPbase^13^. **b**. Phylogenetic tree of FPs constructed from amino acid sequences. **c.** (Left) Scatter plot of FPs by maximum excitation and emission wavelengths based upon reported spectral data for 40 FPs from FPbase^13^). (Right) Spectral distribution represented by determining the full width at 75% of the maximum for each FP’s reported excitation and emission spectra from FPbase^13^. The distributions are marked by intersecting bars (horizontal = excitation, vertical = emission). **(d-e).** Schematic overview of the hyperspectral imaging workflow. **d.** Depiction of the 32-channel spectral detector. Emitted light is simultaneously captured across multiple wavelengths (410*nm* – 695*nm*). Each of the 32 channels collects light in ∼8.9*nm* bins. **e.** Overview of 4-block imaging acquisition. **f.** 100-channel emission spectra acquired from an EGFP monoculture.

To address this barrier, we have developed the Cosine Similarity Decoder of XFP (*CoSiDeX*) toolbox for standardizing FP spectral compatibility. Here we describe how applying a hyperspectral imaging approach with a simple mathematical predictor of FP spectra, allows for accurate detection of spectrally resolvable palettes, from three up to 12 unique FPs. This doubles the previously reported palette of 6 FPs, providing expanded combinatorial co-culture FP-palettes for tracking clonal diversity in living systems. Additionally, we develop an application for importing, segmenting, and re-coloring hyperspectral images by FP identity. Collectively, our work advances the accessibility of hyperspectral imaging and FP reporters thereby laying the foundation for more robust multiplexed lineage tracing in living systems.

## Results

### Hyperspectral Analysis Workflow

A collection of forty-four candidate FPs from multiple species and other synthetic sources (Figure 1b) were selected and cloned from FP base^13^ and used to develop a hyperspectral FP library. Each FP was gene-synthesized in-frame with a 3x-nuclear localization signal and then cloned into a multisite Gateway cloning system^26^ for easy shuttling into eukaryotic expression vectors (Supplemental Figure 1). Each FP is cofactor independent, has a peak excitation and emission in the visible light spectrum (400-700 nm), pKa <6 (except for EYFP, pKA = 6.9), and estimated brightness of > 10 (except for 4FPs, included in our collection for having an Em_max_ > 597nms) (Supplementary Table 1). Although emission and excitation maxima show the spectral diversity of FPs in our library (Figure 1c, Left), the broad range of excitation and emission spectra reduce the likelihood that each FP can be resolved using single or multichannel acquisition, thereby illustrating the need for hyperspectral strategies for combinatorial FP imaging (Figure 1c, Right).

We implemented a hyperspectral imaging approach by performing consecutive scans across 4 different excitation conditions and collecting emission intensity values at 8.9*nm* resolution for each scan. This resulted in four image series of 31 channels (405*nm* excitation), 27 channels (458/561*nm* excitation), 22 channels (488*nm* excitation), and 20 channels (514/594*nm* excitation) totaling 100 channels per region of interest (Figure 1d-e). Each 100-channel stack was maximum intensity projected (MIP) to a single image for CellPose^27^ segmentation. The resulting image masks were applied to each of the 100-channel images, and mean pixel intensity was calculated for each channel. This resulted in a Cell x Channel Intensity matrix for every segmented cell. We tested this workflow using EGFP as a prototype. Human embryonic kidney (H293T) cells were transiently transfected with EGFP and then imaged. Each image produced identifiable EGFP emission spectra whose intensity was dependent upon expected differences of excitation efficiency at each excitation source (Figure 1f, top). The concatenation of each spectrum resulted in a 100-channel EGFP spectrum (Figure 1f, bottom).

### Exploring and quantifying fluorescent protein spectral space

Next, we characterized all 44 FPs in our collection using our hyperspectral imaging and analysis workflow. Each FP was transiently and singly transfected into H293T cells then fixed and imaged at single cell resolution. First, we calculated the integrated brightness of each FP across all 100 channels and found that 37/44 FPs had an average relative intensity between 10^4.5^ and 10^5.5^ while 7/44 of the FPs were more than 2.5 orders of magnitude dimmer (Figure 2a and Supplemental Figure 2). These 7 FPs were excluded from subsequent analyses. Next, we applied a dimensionality reduction analysis of the remaining 37 FP spectral space to determine if subtle differences across each 100-channel spectrum could separate spectrally similar FPs. Uniform manifold approximation and projection (UMAP) and clustering of the 562,534 total single cells (6632 – 21334 cells per FP, average of 12,784 cells/FP) enabled projection and clustering of spectral data (Figure 2b). We previously developed Cancer rainbow mice (Crainbow) that express spectrally resolvable mTFP1, EYFP, and mKO^22^. We found that each of these FPs clustered in their own unique cluster (Clusters 20, 13, and 16 respectively). In contrast, spectrally similar FPs, like Aquamarine, mCerulean3, and mTurquoise2 clustered together (Cluster 1).

**Figure 2.**
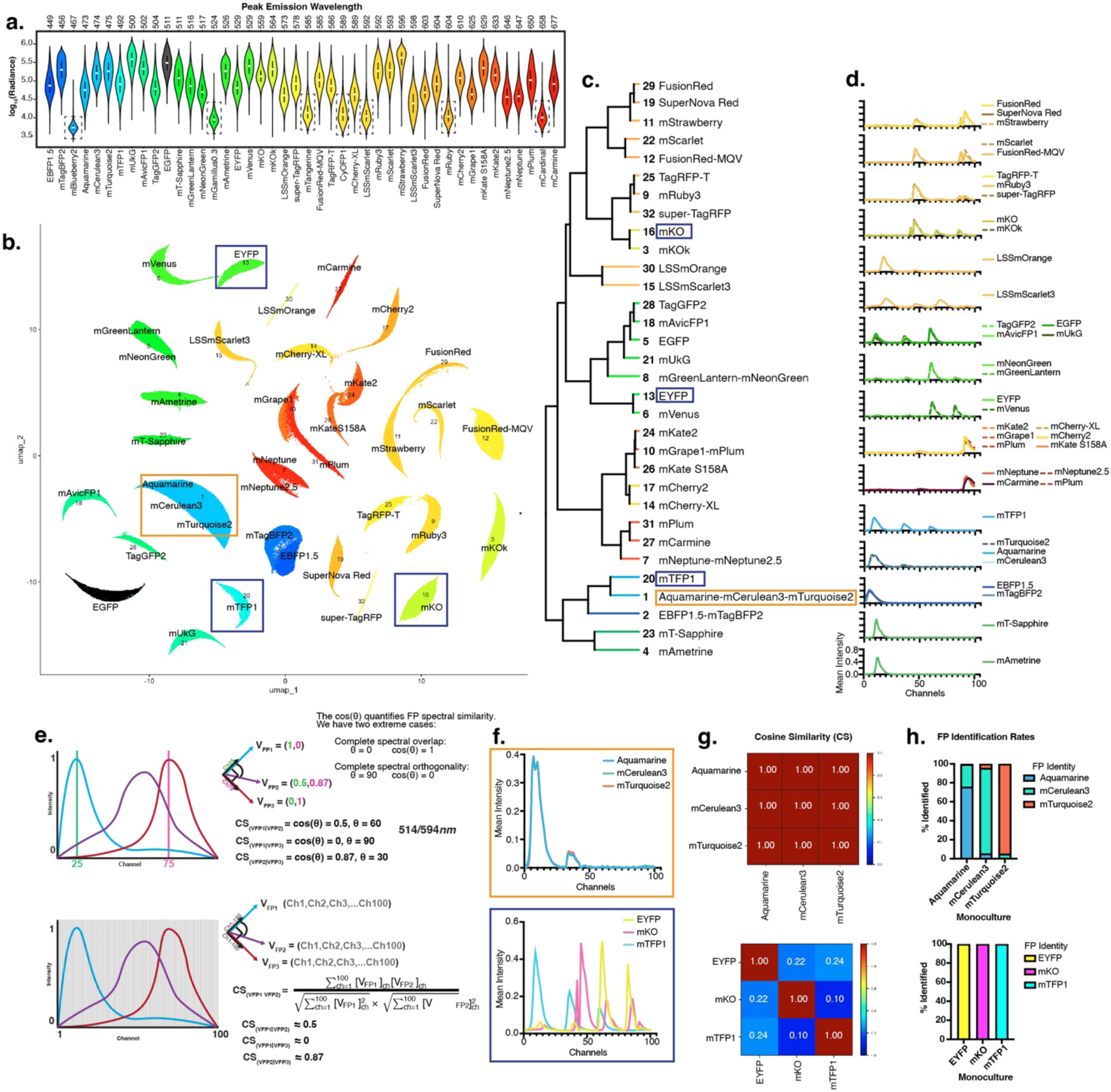
Exploring hyperspectral fluorescent protein space. **a.** Violin Plot showing dynamic range of FP signal intensity. FPs are ordered and colored by wavelength of Em_max_. For each cell within the FP reference monocultures, radiance is defined as the sum of mean pixel intensity across all 100 channels. FPs in black boxes were identified as “low intensity” through hierarchical clustering of pairwise Wasserstein distances between the radiance distributions. (Supplementary Figure 2). **b.** UMAP visualization of FP 100-channel spectra. Cluster number and FPs are overlayed (32 clusters identified). Boxed clusters denote FPs used in panels “f-h”. **c**. Phylogenetic analysis showing hierarchical relationships between UMAP clusters. **d.** Plots showing the reference spectra of FPs within clusters. **e.** Overview of the cosine similarity (CS) calculation when FP spectra are represented as vectors. (Top) In a simple 2D vector space (emission intensity at Ch25 and Ch75), a quantitative measure of emission similarity between FPs is given by CS_(VFP1|VFP2)_ = cos(angle between vectors). CS_(VFP1|VFP2)_ = 0 represents complete orthogonality and CS_(VFP1|VFP2)_ = 1 represents complete overlap. In a 100-dimension vector space, FP spectral similarity is calculated by the represented equation. **f.** Reference spectra of FPs within the same cluster (Top, Cluster 1 FPs, Aquamarine, mTurquoise2, mCerulean3) and FPs from different clusters (Bottom, Crainbow FPs used as an example: EYFP (Cluster 13), mTFP1 (Cluster 20), and mKO (Cluster 16)). **g.** Heatmap showing CS values between FPs. CS of Cluster1 FPs = 1 and CS of Crainbow FPs ranges from 0.1-0.24. **h.** Monoculture FP identification rates using maximum cosine similarity (MCS). For each cell within the FP reference monocultures, CSs were calculated between the cell spectrum and the reference FP spectra of FPs of interest. FP identity was then assigned for each cell based on MCS. For each monoculture, the bar plot shows the % of cells identified as expressing those FPs (Top = Cluster 1 FPs, bottom = Crainbow FPs).

Phylogenetic analysis was performed to visualize hierarchical relationships between FP spectra (Figure 2c). As expected, this analysis confirmed spectral divergence of mTFP1, EYFP, and mKO and the spectral similarity of Aquamarine, mCerulean3, and mTurquoise2. Next, reference spectra were generated for each FP in the dataset by averaging the mean pixel intensity across all cells at each channel (see Methods). FPs within the same clade (i.e. EYFP vs mVenus) had similar reference spectra, whereas between clade spectra were dissimilar (i.e. mTFP1, EYFP, and mKO) (Figure 2d). Overall, the extensive differences between clades and the UMAP visualization together suggested that our hyperspectral imaging approach captures FP spectral diversity in a way that could support enhanced FP multiplexing.

Even though UMAP visualization was useful for presenting spectral data, this type of analysis necessitates large data sets built upon hundreds to thousands of cells. This would preclude analysis of single cells or small clones in an image. Therefore, we sought to develop a process for predicting FP expression at single cell resolution. Cosine similarity is a widely used mathematical method to determine the similarity between two vectors. In our imaging approach, each FP spectrum can be thought of as a vector with 100 dimensions, where each dimension is the intensity of light at that channel. When vectors have equivalent direction, their cosine similarity is 1. Conversely, a cosine similarity of 0 indicates complete orthogonality between two vectors (Figure 2e). As proof-of-concept we first calculated the pairwise cosine similarity for similar spectra (Aquamarine, mCerulean3, and mTurquoise2) and found cosine similarity values of approximately 1.0, indicating, as expected, that these spectra are nearly identical (Figure 2f-g, top). In contrast, pairwise cosine similarity values for mTFP1, EYFP, and mKO ranged from 0.1-0.24, which quantitatively confirm their spectral resolvability (Figure 2f-g, bottom).

Having shown that cosine similarity provides a reasonable quantitative measure of FP spectral similarity, we next tested the feasibility of predicting FP identity using cosine similarity comparisons. We reasoned that the identity of an unknown spectra could be predicted by determining the maximum cosine similarity (MCS) value with known spectra in our library of fluorescent protein references. In other words, if three FPs are used in an experiment, every cell will be labelled according to its MCS with the three candidate reference spectra. As a test data set, we utilized the previously acquired spectra of individual cells in our reference monocultures (Figure 2a-b). First, we quantified cosine similarity values between each Aquamarine, mCerulean3, and mTurqoise2 expressing cells and the three reference spectra. Then the MCS was used to predict the FP identity. Comparing predicted identity to ground-truth showed that Aquamarine, mCerulean3, and mTurquoise2 expressing cells were susceptible to misidentification using MCS prediction. (Figure 2h top). Next, the cosine similarity of every individual mTFP1, EYFP, and mKO expressing cell was pairwise calculated with all three reference spectra generated for each of these FPs (Figure 2h bottom). We found that MCS accurately predicted all EYFP, mTFP1, and mKO cells in the dataset.

### Developing a <u>Co</u>sine <u>Si</u>milarity <u>De</u>coder of <u>X</u>FP (CoSiDeX) tool for FP identification and visualization

Our next goal was to empirically determine cosine similarity cutoffs for accurate FP identification. Green FPs (TagGFP2, mGreenLantern, mNeonGreen, mT-Sapphire, mUkG, mAvicFP1, and EGFP) were used as proof-of-concept. We plotted reference spectra for these FPs and found that despite overall similarities in Em_max_, three qualitative patterns were observable (Figure 3a). Cosine similarity scores between the FP reference spectra were calculated for each pair of FPs and ranged from 0.15 to 1.00 (Figure 3b). Unsurprisingly, the long-stokes shift FP, mT-Sapphire, is highly orthogonal to the other green FPs as evidenced by low range of cosine similarity values with the other green FPs (0.15-0.58). In contrast the green FPs within pattern 1 (EGFP, TagGFP2, mUkG, and mAvicFP1) are nearly identical to one another with cosine similarity values ranging from 0.95-0.99. Similarly, the green FPs within pattern 2 (mNeonGreen and mGreenLantern) were nearly identical with a cosine similarity of 1.00. Interestingly, the pairwise comparisons of pattern 1 to pattern 2 yield cosine similarity values of 0.81 to 0.87, suggesting that FPs with spectra having a cosine similarity of < 0.9 can be accurately resolved.

**Figure 3.**
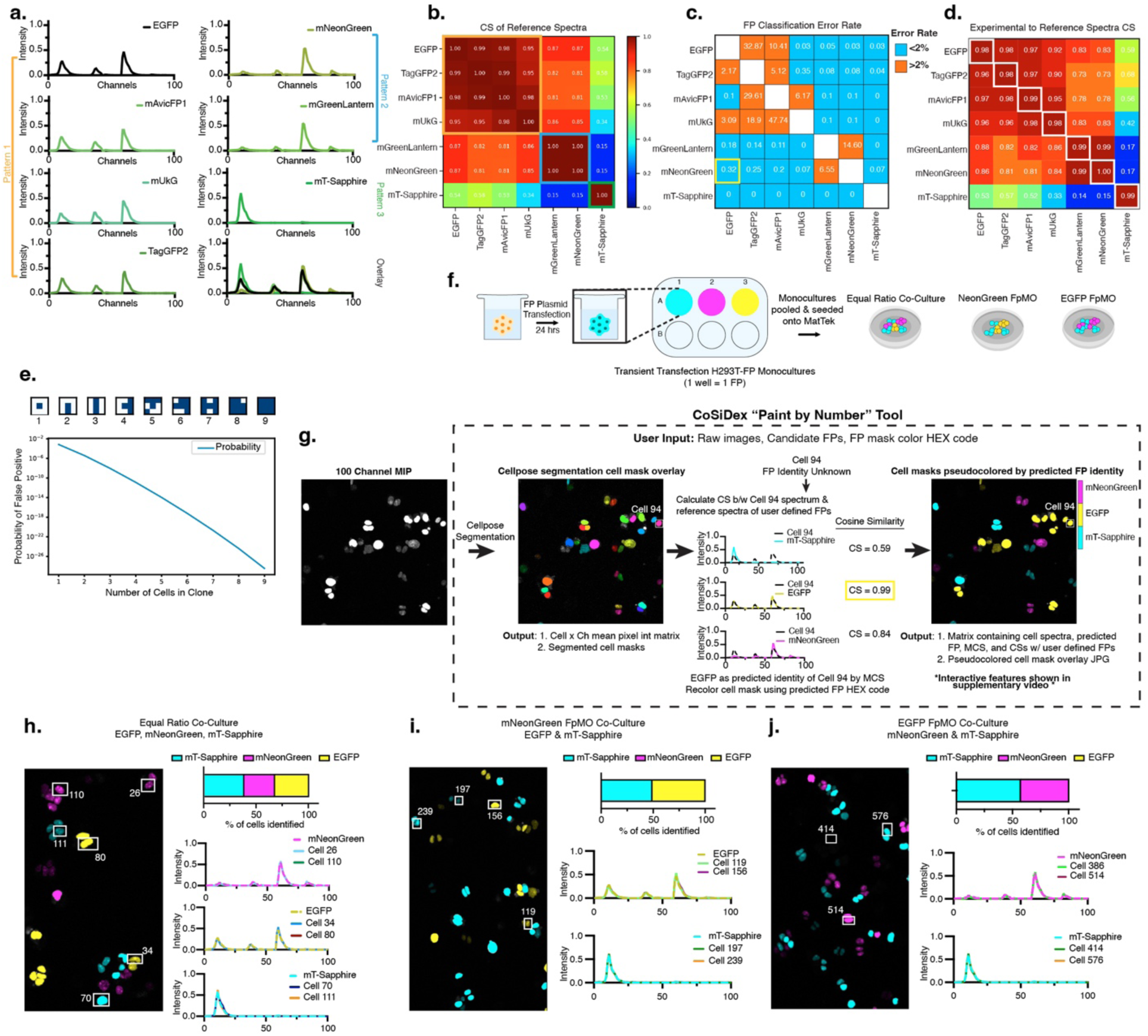
Cosine similarity quantifies FP spectral resolvability. **a.** Reference spectra for green FPs demonstrate three qualitatively distinct spectral patterns. **b.** Heatmap of pairwise cosine similarity (CS) values among green FP reference spectra. FPs within the same pattern exhibit high similarity (Pattern 1: 0.95 < CS < 0.99; Pattern 2: CS = 1.00), whereas comparisons across patterns yield lower CS values (range: 0.15–0.87). **c.** Classification error rate matrix. For each new monoculture, error rates were calculated by performing pairwise comparisons of FP predicted identity when given the choice between the actual FP and every other green FP. An error rate threshold of 2.00% was designated as acceptable (blue: <2.00%; orange: >2.00%). The yellow outline highlights the pair with the highest CS below this threshold (mNeonGreen vs. EGFP; CS = 0.86; error rate = 0.32%). **d.** CS values between new experimental monocultures and FP reference spectra. Transiently transfected H293T cells were imaged for each FP, and CS values between each cell spectrum and all reference spectra were averaged. **e.** %False positive in simulated spatial clone of EGFP cells. Clones of up to 9 cells were simulated in a 3×3 grid (central position fixed), with remaining positions randomized. %False-positive classifications decreased exponentially with increasing clone size (see Methods). **f.** Co-culture experimental design. Monocultures of transiently transfected H293T-FP cells were verified and then reseeded into various co-culture conditions: equal FP ratio, mNeonGreen FP-minus-one (FpMO), and EGFP FpMO. **g.** Overview of *CoSiDeX Paint-by-Number* tool. Hyperspectral imaging generates a grayscale maximum intensity projection (MIP). After segmentation and spectral extraction, FP identity is predicted using maximum cosine similarity. Cells are then pseudocolored using user-defined HEX codes based on their predicted FP identity. **(h-j).** Green FP co-culture analyses. Representative pseudocolored images of co-cultures are shown. For each co-culture, the percentage of cells classified as expressing each FP is plotted. Spectra from representative cells (highlighted with boxes) and their predicted FP reference spectra are plotted on the right.

To assess whether a cosine similarity value of 0.9 is a reliable threshold of spectral resolvability, we performed a classification experiment on newly generated green FP monocultures to evaluate whether MCS could correctly predict FP identity given a binary choice between the correct FP reference and another green FP reference. The percentage of cells misclassified 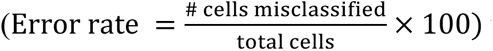 was then calculated for every pairwise FP comparison (Figure 3c). Pairwise comparisons of FPs within the same pattern (i.e. either pattern 1 or pattern 2) had error rates as high as 47.74% (mUkG vs. mAvicFP1). This indicates that if mUkG and mAvicFP1 were used in the same experiment, then approximately 47% of the true mUkG cells would be misclassified as mAvicFP1. In contrast, pairwise comparisons between pattern 1, 2, and 3 revealed error rates <2.00% and often 0.00%. These results confirmed that spectra with cosine similarity values as high as 0.87 (mNeonGreen vs. EGFP) can still be accurately discriminated with only a 0.32% error rate (Figure 3c, yellow box). In other words, if 10,000 mNeonGreen and EGFP cells were imaged, approximately 32 cells would be mislabeled.

To determine whether the reproducibility of a FP’s spectra could be impacting error rates, we averaged the cosine similarity values across the experimental monoculture cells and each green FP reference spectra. Pairwise comparisons of the experimental FP spectra to its corresponding reference spectra revealed cosine similarity values approximately equal to 0.98-1.00, indicating that each spectrum is reproducible from one experiment to the next (Figure 3d, white boxes). Furthermore, we modeled probabilistic spatial constraints for assessing error rates for cells within a clone between EGFP and mNeonGreen, from 2 cells up to 9 cells (see methods). If a single EGFP cell existed in a field of 10,000 cells, then our error rate of 0.32% would preclude accurate FP labeling because 32 cells would be misidentified. However, if 2-3 adjacent cells are named EGFP then our error rate is approximately 1 mislabeled cell per 1,000,000 counted cells. As the number of neighboring cells grows larger, this error rate continues to fall significantly (Figure 3e).

Next, we solved for the best spectrally resolvable green FPs for use in multiplexed imaging. We solved for all spectrally resolvable combinations (“N-tuple”) of green FPs from 2-tuples up to 7-tuples by first excluding any combination where pairwise cosine similarity values were greater than or equal to 0.9. We found 13 possible 2-tuples and 8 possible 3-tuples (Supplementary Figure 3). No combinations of 4- to 7-tuples were found. Next, we performed validation experiments to directly test the resolvability of a representative 3-tuple (mT-Sapphire, mNeonGreen, and EGFP) (Figure 3f). Cells transiently transfected with either mT-Sapphire, mNeonGreen, or EGFP were mixed in a single co-culture (1:1:1), allowed to grow for 2 days and then fixed and imaged. The MCS for each cell was calculated using mT-Sapphire, mNeonGreen, and EGFP reference spectra and the cell masks were recolored according to each predicted FP using the *CoSiDeX Paint-by-Number* tool (Figure 3g, Supplementary Video 1). Qualitatively, most FP labels contained at least 2 cells within a spatial clone, increasing the likelihood that each cell was named correctly as modeled in Figure 2E. On a population level, we observed the expected proportion of cells expressing each FP in an equal mix (Figure 3h). We also visually confirmed that experimental cell spectra matched the predicted reference spectrum (Figure 3h). To further verify accuracy of FP predictions, we also performed fluorescent protein minus one (FpMO) controls. Cells mixed 1:1 (mT-Sapphire and EGFP) were imaged, and MCS predictions were made using mT-Sapphire, EGFP, and mNeonGreen reference spectra. EGFP and mT-Sapphire spectra were retrievable and as expected, the culture was labelled approximately 48.74% mT-Sapphire, 51.16% EGFP, and 0.10% mNeonGreen (Figure 3i). Similar results were found with a mT-Sapphire and mNeonGreen mix (57.07% and 42.93% respectively, Figure 3j).

A substantial concern with spectral unmixing is the intensity dependent contributions to unmixing between experiments and adjacent spectra. Cosine similarity uses normalized spectra such that the shape and not the intensity is directly compared. To determine whether intensity affects the accuracy of FP naming, we divided cells in the mT-sapphire, mNeonGreen, and EGFP monocultures, into low, medium, and high intensity groups. In each case, the FP classification error rates were 0% except for mNeonGreen which had an error rate of 0.33% in the low intensity group (Supplementary Figure 4). Altogether, this demonstrates we can resolve multiple green FPs using MCS, providing a rationale for applying similar strategies across all FPs used.

### Using CoSiDeX for rationally selecting FPs for combinatorial experimental biology

As previously mentioned, a major limitation in FP-based tool development is how to rationally choose the most compatible FPs for an experiment. To test the ability of our approach to predict scalable combinatorial FP palettes across the visible light spectrum, we returned to our reference data set and calculated pairwise cosine similarity values (37 FPs, Figure 4a). Next, we systematically calculated all possible FP N-Tuple palettes from (N = 2 to N = 12). The N-tuple palettes with the lowest possible MCSs, and therefore predicted to be the most resolvable, are shown for N = 4-12 FPs (Figure 4b).

**Figure 4.**
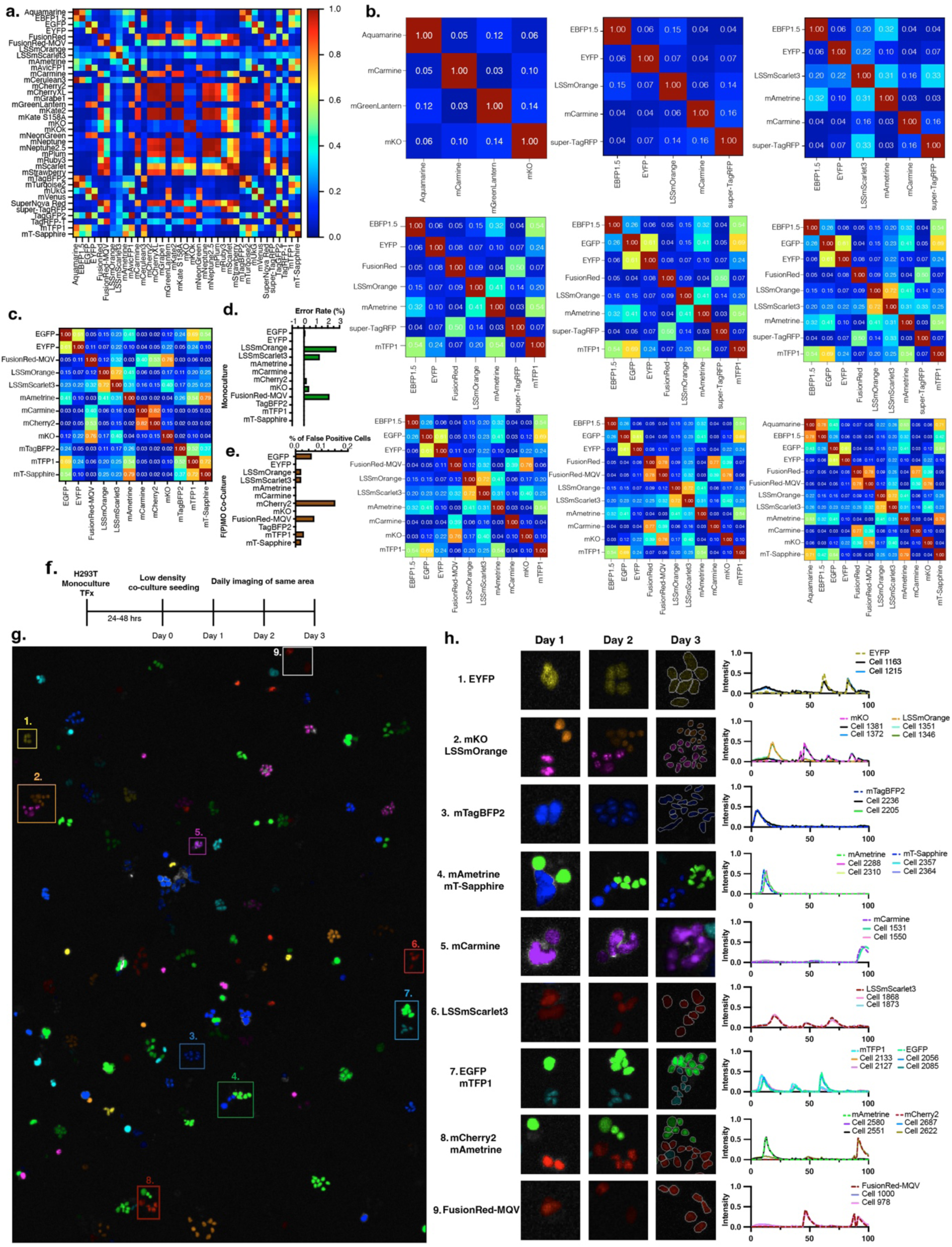
Tracing growth of 12 unique FP clones using the CoSiDeX Toolbox. **a.** Heatmap of pairwise CS values for FPs in our library. **b.** Heatmaps of pairwise CS values of optimized FP palettes ranging from 4- to 12-Tuples. **c.** CS heatmap for 12-Tuple combination compatible with Crainbow FPs. **d.** FP identity prediction error rate in new transient transfection monocultures when given the choice of all 12 FPs (Max error rate = 2.2%). **e.** %False positive classifications in 12-Plex FpMO co-cultures when given the choice of all 12 FPs (Max %False Positive = 0.15%). **f.** Schematic of 12-Plex clonal growth experiment. **g.** Day 2 image from 12-Plex clonal growth experiment pseudocolored cell masks by predicted FP (Tracked clones outlined by white boxes). **h.** Example images of clonal growth tracking. For each FP, the same colony is shown on Day 1 – 3 (Left). Outlines of cell masks added in Day 3 for dim cells. Example spectra of cells within each colony at Day 3 plotted along with the predicted FP reference spectra (Right).

Previous work has demonstrated resolvability of up to 10 FPs using a hyperspectral flowcytometry approach. Therefore, to assess the power and accuracy of our N-Tuple predictions for microscopy, we experimentally validated a predicted 12-Tuple FP palette (Figure 4c). First, new experimental FP monocultures were generated, and FP classification error rates were calculated when given the choice of all 12 possible FP reference spectra. All but three monocultures had error rates of <0.05%, and the remaining three monocultures had error rates ranging between 1.30% and 2.71% (average error rate = 0.57%, Figure 4d). Next, we conducted a false positive analysis to determine whether or not 12 different FP-tagged cell lines could be imaged in the same sample. Fluorescent protein minus one (FpMO) co-culture conditions were generated for each of the 12 FPs. Quantification across FpMOs demonstrates that our approach results in almost undetectable proportions of false positives (%False Positive range of 0 – 0.15%, Figure 4e). Combined, these experimentally determined error rates and false positive detection proportions suggest that our method’s FP prediction reflects reliable spectral distinctions with low spectral artifacts and enables multiplexed fluorescent protein imaging.

A potential application of this expanded FP resolvability could be tracing growth dynamics/interactions of 12 distinct cell types, with each FP barcoding a unique biological marker. Given that our co-culture experiments have relied on proportions of fixed cells, we next sought to determine whether the same population of cells could be consistently traced over time in a spatially relevant context. Briefly, 12 FP monocultures were generated by transient transfection and mixed into a co-culture seeded at a low density. To track clonal expansion, the same area of the plate was imaged daily for 3 days (Figure 4f, Supplementary Figure 5). Examples of a representative clonal patch tracked across each day are shown for each FP (Figure 4g, 4h Left). Spectra of representative cells and their predicted FP within the day 3 clonal patches are shown (Figure 4H right). Of note, the intensity of the FP signal decreases across the days. This is consistent with the dilution of plasmid across cell divisions that is expected in transient transfections. This provides further support for both the fact that we are tracing biologically related clones, and that FP classification is not impacted by brightness. Altogether, these experiments demonstrate reliable prediction of up to 12 unique FPs and one application for tracing clonal behavior in living cell.

Having validated the resolvability of our N-Tuple FP predictions, we have developed the *CoSiDeX Palette Generator* tool for predicting N-Tuple palettes. Using our FP reference spectra, this tool generates potential FP palettes using either user defined FPs to include or exclude (N = 2-15 Tuples). The *Palette Generator* creates cosine similarity heatmaps of optimized or user selected FPs, and plots the selected FP’s reference spectra (Supplementary Video 2). Additionally, while the reference spectra for our FP library comes pre-loaded, users can load their own set of reference spectra into the tool for added flexibility. Overall, this tool serves as an objective resource for quantitatively predicting FP resolvability and democratizing hyperspectral imaging analyses.

## Discussion

Here, we describe *CoSiDeX*, a resource and computational framework for expanding FP multiplexing to extend parallel measurements beyond the conventional two- or three-color systems. Our approach combines hyperspectral imaging and cosine similarity, to facilitate a fine-grained, quantitative assessment of FP spectral resolvability. This addresses a major challenge shared by most experimental biologists who seek to reliably image dynamic and complex biological systems. As part of the *CoSiDeX Toolbox*, we provide a suite of tools that streamline FP utilization. To aid with experimental planning, we provide the *CoSiDex Palette Generator* which utilizes <u>co</u>sine <u>si</u>milarity to predict N-Tuple palettes based on user-defined selection criteria. On the analysis end, we provide the *CoSiDeX Paint-by-Number* tool that segments cells, <u>de</u>codes FP identity using MCS, and then pseudocolors cells accordingly.

Our workflow begins with a hyperspectral imaging approach that iteratively acquires emission data for various excitation sources. This yields ∼100 channels of measurements that when concatenated, maximize resolvability between FPs. Similar imaging workflows have been described previously for other fluorophores^28^, that together with our work demonstrates the importance of maximizing excitation diversity for optimal spectral resolution. Spectral data are then represented in vector space for cosine similarity calculations. This approach has previously been utilized to assess spectra stability in hyperspectral flow cytometry datasets^29^ and in spatial transcriptomics to resolve transcriptional states^30,31^. Strikingly, both our data and the transcriptomic analyses indicate that a cosine similarity threshold of 0.9 is sufficient for reliably resolving distinct FPs or cell types.

Resources for choosing the best FPs to use in flowcytometry^32,33^, predict but do not validate, a palette of 10 FPs^32^. We show that by calculating combinations that preserve the lowest possible cosine similarity we were able to solve for the best FPs to use across N-tuples for 3 FPs and up to 12 FPs. This solves a significant problem faced by most biologists seeking to apply FPs to experiments requiring parallel tagging of multiple processes.

The cosine similarity approach diverges from traditional linear unmixing in a few ways. Linear unmixing deconvolves spectral data by finding the optimal linear combination of component spectra (i.e. user-specified references) that make up the target spectrum being analyzed^34^. This computationally intensive analysis results in a set of non-negative weights that specify the strength of each component spectrum. This means linear unmixing is sensitive to signal intensity fluctuation. In contrast, our <u>co</u>sine <u>si</u>milarity-based <u>de</u>coding approach finds the single spectrum that correlates most strongly, across all channels, with the unknown spectrum. This pairwise spectral comparison is trivially parallelizable, substantially improving efficiency for large datasets. However, because the capacity of cosine similarity to resolve combinatorial signatures within individual cells remains an open question, linear unmixing remains the best option for combinatorial analysis.

We demonstrate the practical use of the *CoSiDeX Toolbox* in a proof-of-concept clonal tracing experiment where we show simultaneous visualization of 12 distinct somatic clones in live imaging. Such approaches have broad utility for tracking cell lineage dynamics and clonal competition in models of development, tissue regeneration, and malignancy. Our data also provide clarity on the spectral diversity of FPs within a species. For instance, we predict that 7/11 FPs from *Aequorea Victoria* can be resolved (Supplementary Figure 6). Leveraging phylogenetic similarity in genetically modified organisms such as the immunologically tolerant GFP mice^35^, could prove useful in addressing potential effects of immunogenicity to FPs while still enabling combinatorial FP imaging.

Overall, our work provides a resource of empirically validated and ready-to-use FP plasmids for hyperspectral imaging, along with the *CoSiDeX Toolbox.* Our work serves as a foundational resource for advancing next generation FP-based molecular tools for dynamic and spatiotemporal analysis across diverse biological systems.

## Materials & Methods

### Fluorescent protein selection

FPs were selected from FPbase based on the following criteria: pKa < 6, brightness > 20 (relative scale), cofactor independence, and representation across multiple species. Initial candidates were filtered by emission spectra spanning 400–700 nm and categorized into seven spectral classes: blue (420–470 nm), cyan (470–500 nm), green (500–525 nm), yellow (525–555 nm), orange (555–580 nm), red (580–630 nm), and far-red (630–700 nm). Within each class, FPs were selected to maximize spectral separation while favoring lower pKa values and broader phylogenetic origins when multiple options exhibited similar emission profiles. For example, mVenus was selected over mGold despite slightly lower brightness (66.56 vs. 68.48) due to its more favorable pKa (5.5 vs. 5.9). In spectral regions where FP diversity was limited— particularly red and far-red—brightness thresholds were relaxed to ∼10. Three exceptions were included based on spectral uniqueness despite low brightness: mGrape1 (1.5), mPlum (4.1), and mCarmine (5.81). Complete FP properties are listed in Supplementary Table 1.

### Fluorescent protein cloning

Plasmid designs and Gateway cloning^26^ were performed *in-silico* using SnapGene software (www.snapgene.com). Double stranded FP DNA fragments were synthesized as gBlocks from IDT (Coralville, IA, USA) based on amino acid sequences from FPbase^13^. FPs were flanked with attB recombination sites compatible with either two- or three-fragment Gateway cloning and modified to include a 5’ Kozak sequence + triple nuclear localization signal (5’-gccaccatggacccaaaaaagaagaggaaggtggaccctaaaaagaagcgaaaagtcgatcccaagaaaaaaaggaaggtt-3’), as well as a 3’ universal primer site (5’-gttcaccttttcgaatcctcg-3’).

For two-fragment Gateway reactions, attB1-FP-attB2 fragments were recombined into pDONR221 and pcDNA3.1-DEST (Thermo Fisher Scientific) vectors using a one-step BP:LR reaction (25:75 ratio)^36^. Reaction mixtures contained 25–30 ng FP gBlock, 150 ng each of pDONR221 and pcDNA3.1-DEST, 1 μL BP Clonase II, 2 μL LR Clonase II (Thermo Fisher Scientific), and 2 μL TE buffer. Reactions were incubated at 25°C for 6 h, terminated with Proteinase K and transformed into Mach1 chemically competent cells (Thermo Fisher Scientific, C869601). Following recovery in SOC medium, transformations were plated onto LB agar supplemented with kanamycin or ampicillin. Single colonies were expanded overnight at 37°C, and plasmid DNA was purified using the QIAprep Spin Miniprep Kit (QIAGEN). Constructs were validated by restriction enzyme digestion and Sanger sequencing (Eton Biosciences, San Diego, CA, USA).

For three-fragment Gateway reactions, attB4-FP-attB3 fragments were cloned into pDONR P4r-P3r vectors, and final constructs were assembled into PiggyBac transposon backbone, a gift from Christopher Newgard (Addgene plasmid #121875), according to the Gateway Cloning Technology manual or as previously described³⁵.

### Cell Culture and FP monoculture generation

HEK293T cells (ATCC CRL-3216) were cultured in high-glucose DMEM supplemented with 10% fetal bovine serum and 1% penicillin–streptomycin and maintained at 37°C in a humidified incubator with 5% CO₂. Cells were seeded either onto tissue culture-treated plasticware or poly-D-lysine (PDL)-coated glass-bottom dishes (MatTek Life Sciences) depending on experimental requirements. Transient transfection was performed using calcium phosphate precipitation. Briefly, HEK293T cells were seeded at uniform confluency (50-70%). After 24 hours, each well was transiently transfected with a single FP-encoding plasmid from our mammalian expression library using a calcium phosphate transfection method. For monoculture experiments, transfections were performed directly into PDL-coated MatTeks, fixed with 10% neutral-buffered formalin 1-2 days after transfection, and stored in phosphate-buffered saline until imaging. For co-culture assays, FP-expressing monocultures were generated by transient transfection and imaged 24 h post-transfection to assess transfection efficiency. Cells were then dissociated with trypsin and re-seeded into PDL-coated MatTek dishes. Co-cultures were fixed between 1–4 days post-seeding, depending on experimental design.

### Phylogenetic analysis of FP species

Amino acid sequences of FPs in a FASTA format were aligned using MUSCLE (msa package^37^, R). Pairwise sequence similarity distances were calculated (seqinr packages^38^), and hierarchical clustering was performed. The resulting dendrogram was converted to a phylogenetic tree (ape package^39^) and visualized using ggtree^40^. Final figures were exported as PDFs.

### Hyperspectral imaging

All imaging was performed on a Zeiss LSM 880 laser scanning confocal microscope (Carl Zeiss, Thornwood, NY, USA) equipped with a 32-channel GaAsP spectral detector and seven laser lines. Spectral images were acquired using a 10X objective and an 8.9 nm channel bandwidth in lambda mode. Imaging conditions were held constant across experiments unless otherwise specified (e.g., tiling or z-stack parameters; see Supplementary Table 2 and Figure 1d).

Four sequential spectral acquisition blocks were collected per field of view: Block 1 (405*nm* laser, Emission 420-695*nms*), Block 2 (458/561*nm* lasers, Emission 455-695*nms*), Block 3 (488*nm* laser, Emission 500-695*nms*), and Block 4 (514/594*nm* lasers, Emission 517-695*nms*). This results in 4 separate image files per sample. For live imaging experiments, MatTek dishes were placed in a live-imaging stage-top incubation chamber (PeCon GmBH, Erbach, Germany).

### Cell segmentation and spectra generation

Following hyperspectral image acquisition, the 4 block image files were combined sequentially into a single 100-channel stack. The stack was then maximum intensity projected (MIP), to create a single image. Segmentation was performed on the MIP using a fine-tuned Cellpose 2.0^27^ pipeline, resulting in masks for each cell. Cell masks were then applied to each channel of the image stack, and the mean pixel intensity was calculated at each channel for each cell mask. This resulted in a Cell x Channel Intensity matrix (n x 100), with rows corresponding to n cells and columns corresponding to the channels. Reference spectra for the 44 FPs in our library were generated in parallel from transient transfection monocultures, seeded into a 48-well glass bottom imaging plate (MatTek, P48G-1.5-6-F) as described above. All wells were imaged sequentially in a single imaging session using the hyperspectral approach described above. Spectral matrices for each well were generated as detailed above. For each monoculture, spectral information was obtained from between 6632 – 21334 cells (average of 12,784 cells/FP).

### FP relative brightness

To quantify the signal intensity of FPs expressed in mammalian cells, radiance was calculated from the reference monocultures spectral matrices by summing the mean pixel intensity across all 100 channels. To normalize distributions for comparison, a base-10 logarithmic transformation to the per-cell radiance values was performed and visualized as violin plots. FPs were color-coded and arranged by ascending peak emission wavelength, using published spectral data from FPbase^13^. Clusters of FPs with similar brightness profiles were identified using pairwise Wasserstein distances followed by hierarchical clustering (SciPy^41^). This revealed a distinct cluster of 7 ‘dim’ FPs with similar radiance characteristics that were lower than the remaining 37 FPs.

### UMAP/Clustering

Spectral matrices from each FP reference monoculture were concatenated into a single matrix and imported into R (version 4.2.3). The seven ‘dim’ FPs (mCardinal, mBlueberry2, CyOFP1, Gamillus0.3, mTangerine, LSSmScarlet, mRuby) were filtered out of the matrix before creation of a seurat object and normalizing the data using Seurat (V5.2.1)^42^. The expression matrix was transposed to scale the data by cell instead of channel in order to cluster based on variance in spectral signal not brightness. Principal components analysis was performed, and elbow plot was used to find the number of PCs that describe the variance to *FindNeighbors* and *RunUMAP*. Cluster resolution was determined to be 0.4 by choosing the resolution that most closely matched the number of FPs imaged. Cluster tree was built using the *BuildClusterTree* function.

### Cosine similarity

Treating FP spectra as a 100-dimension vector, for any two vectors *V_FPa_, V_FPb_*, the cosine similarity is defined as 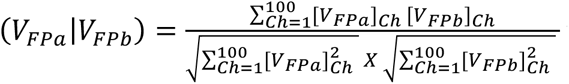 where [*V_FPa_*]*_ch_* and [*V_FPb_*]*_ch_* denote the mean pixel intensity of the vectors at a given channel (Ch). Here, the numerator computes the scalar product of the two vectors, capturing their alignment, while the denominator normalizes the result to 1 by calculating the product of their Euclidian norms. This normalization ensures that comparisons reflect differences in spectral shape rather than absolute intensity, with 1 indicating identical vectors and 0 indicating no correlation.

### Predicting FP Identity, CoSiDeX Paint-by-Number Tool

To predict the FP identity of an unknown spectrum, i.e. to classify a given cell by its spectrum, cosine similarity values between the unknown spectrum and each reference spectrum within a predefined set FPs were calculated. The FP with which the unknown spectrum had the maximum cosine similarity (MCS), was predicted to be the FP expressed by that cell. For any given sample, this was performed on a per cell basis for all segmented cells. To assess accuracy of FP prediction, two separate calculations were utilized. In the setting of a monoculture experiment, where the FP identity is known, we used 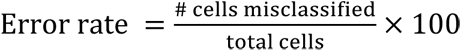, to quantify the % of cells misclassified in the monoculture. In FpMO co-culture experiments, where the FP identity for any given cell is truly unknown, we utilized the ground truth of expecting 0 cells from the missing FP to assess accuracy of classification. In this case, we calculated misclassification as 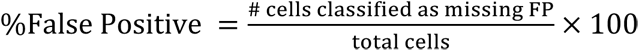

To further aid analysis, we designed the CoSiDeX *Paint-by-Number* application. This is a custom python application developed to integrate the Cellpose^27^ segmentation and spectral classification for visualization and exploratory analysis. After segmentation, spectral generation and FP identity prediction using MCS (detailed above), cell masks are pseudocolored based on their predicted FP identity. Each FP is assigned a RGB color code which can be specified by the user. Cell mask pseudocoloring is performed on the original MIP, with cell mask intensity scaled to reflect cell radiance, while non-cell regions are rendered in grayscale to maintain visual clarity. Through an interactive interface, application also includes several modular features designed to support analysis: (i) the option to toggle cell outlines to enhance visibility; (ii) a global color toggle to disable pseudo-coloring; (iii) a color bar displaying counts and labels of each FP type, which can be used to filter displayed cells by clicking on the color of interest; and (iv) the ability to click individual cells to display their spectral profile, with selected cells highlighted using a complementary outline color.

### N-Tuple Prediction, CoSiDeX Palette Generator

The optimal N-tuple palette of FPs from a library, were defined as those which minimize the sum of all cosine similarity values among pairs of candidates. As part of the *CoSiDeX* Toolbox, we have created an app that generates optimal FP palettes from 2- to up to 15-tuple combinations. To predict optimized N-tuple FP palettes, we first eliminate FPs with cosine similarity values > 0.9 from the list of options. Therefore, if any two spectra have such a high similarity, only one is kept. This pre-filtering step reduces the space of possible solutions making the search more efficient. Next, all pair-wise cosine similarity values among remaining FPs are calculated. Then, all possible N-tuple combinations are generated and all cosine similarity products among the pairs of candidates are summed. The N-tuple combination which results in the minimum sum of cosine similarity values is selected as the optimized FP palette. The *CoSiDeX FP Palette Generator* app comes preloaded with our 37 FP reference spectra, but users can load their own reference spectra file. The user can then define the number of FPs desired in the palette, and any FPs requirements (both to include or exclude). The optimal palette is then calculated, and the user can then select those FPs to generate a heatmap of cosine similarity values, which can be exported as a .png. To aid with visualization of spectral overlap, an additional feature of the app is the ability to plot the spectra of any selected FPs.

### Spatial Clone Analysis

The impact of spatial distribution on FP misclassification was assessed through probabilistic modeling. Spatial distribution of a clone of N EGFP cells (up to 9) in a 3×3 grid were simulated with the center square always filled, and the remaining N-1 cells placed randomly. For a spatial clone of N EGFP cells, there were 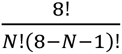 ways of placing them, and a probability of *P^N^* for those cells being misclassified as mNeonGreen.

## Supporting information

CoSiDeX_SupplementaryFigures

## Code Availability

Code will be made to reviewers and editors upon request and will be made publicly available upon publication.

## Acknowledgments

The authors would like to thank Dr. Talley Lambert for kindly sharing code to query Fpbase.org. This study was funded by a grant to JCS and HKL (NCI R21CA267012).

## Author Contributions

J.D.G., H.K.L., and J.C.S. conceived the project. A.V.P. and J.C.S. designed the experiments. A.V.P. and J.E.S. performed experiments, acquired data, and cloned plasmids. A.V.P., H.C., J.D.G., B.W.R., and J.E.D. analyzed data. H.C. and J.E.D. developed software, with input from A.V.P. A.V.P. and J.C.S. wrote the manuscript. All authors reviewed and edited the manuscript. J.C.S. supervised the entire project.

