## Supplementary material for "CoSiDeX: A hyperspectral fluorescent protein resource for highly multiplexed imaging": CoSiDeX_SupplementaryFigures: CoSiDeX_SupplementalFigures.pdf

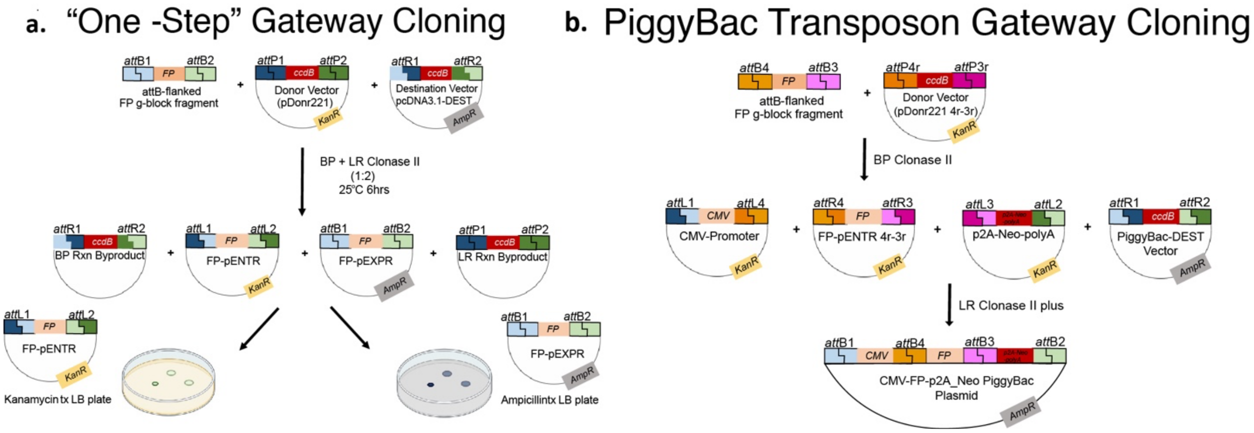


**Supplementary Figure 1. FP plasmid generation. a.** Overview of the "One-step" BP + LR Clonase II gateway cloning protocol for pENTR and pEXPR FP plasmid generation. All reagents for both BP and LR reactions are combined in a single tube for simultaneous reaction performance. FP-pENTR and FP-pEXPR plasmids are selected for by plating on appropriate antibiotic treated agar plates. **b.** Overview of FP-PiggyBac plasmid generation. FP-pENTR plasmid is generated via a BP clonase II reaction with IDT synthesized *attB4/B3* FP genes and a pDonr2214r-3r plasmid. A 3-Fragment LR Clonase plus reaction is then performed between a CMV promoter, FP-pENTR, p2A-Neo-polyA, and a PiggyBac-DEST plasmid.

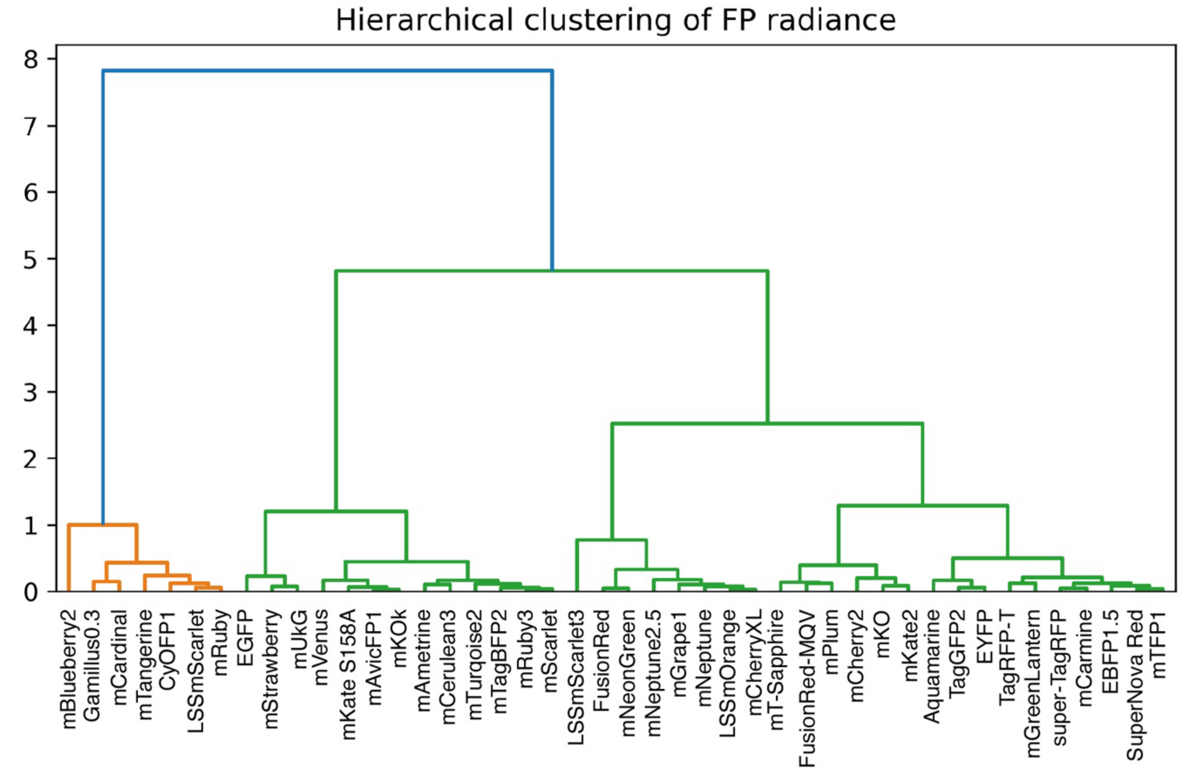


**Supplementary Figure 2. Identification of "low intensity" FPs**. Pairwise Wasserstein distances between FP radiance distributions was calculated followed by hierarchical clustering. The 7FPs clustered on the left (mBlueberry2, Gamillus 0.3, mCardinal, mTangerine, CyOFP1, LSSmScarlet, and mRuby) were identified as "low intensity" and excluded from our library.

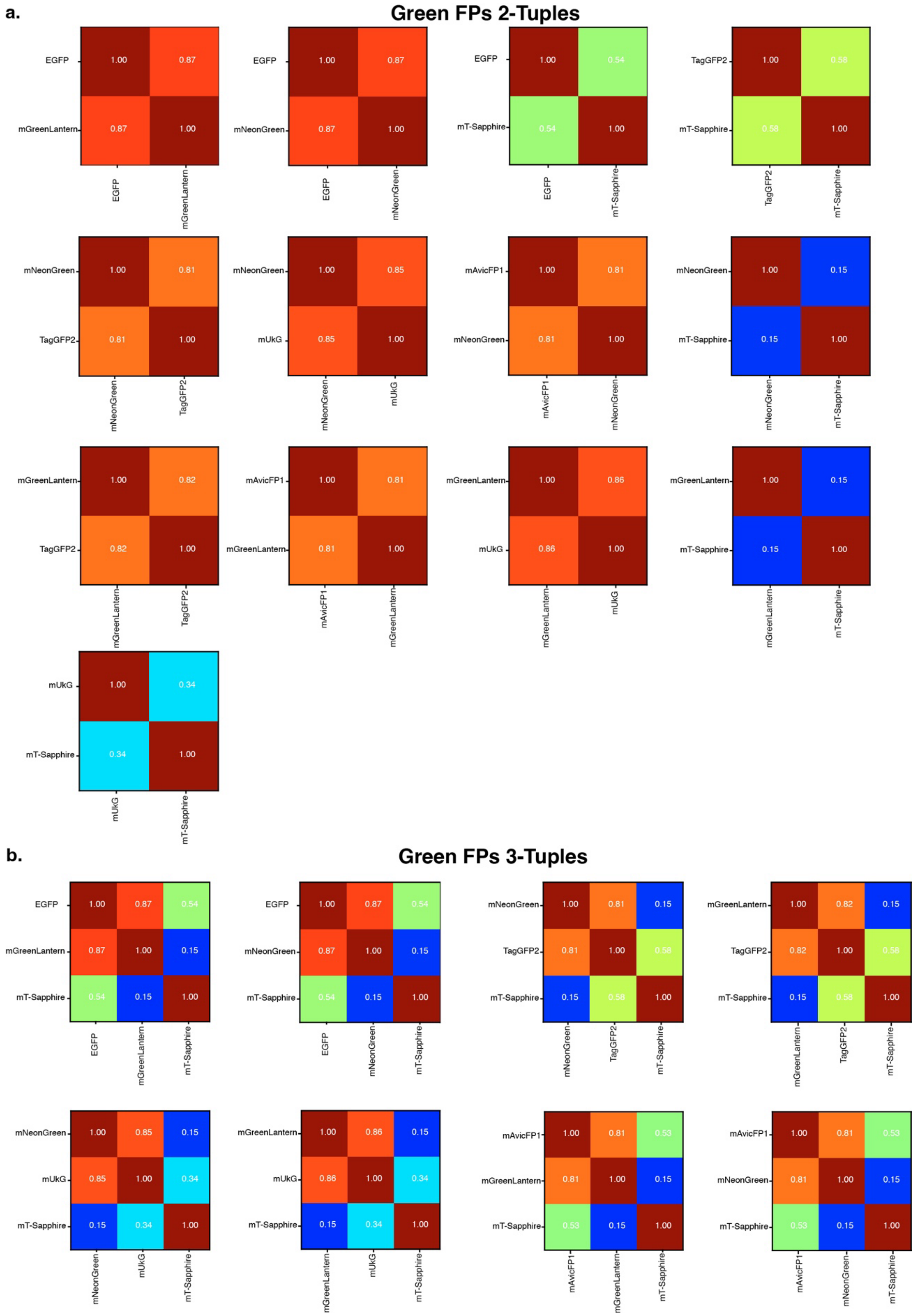


**Supplementary Figure 3. Green FP N-Tuple Palette Predictions. a.** Heatmaps for each predicted resolvable (**a)** 2-Tuple and **(b)** 3-Tuple FP palettes.

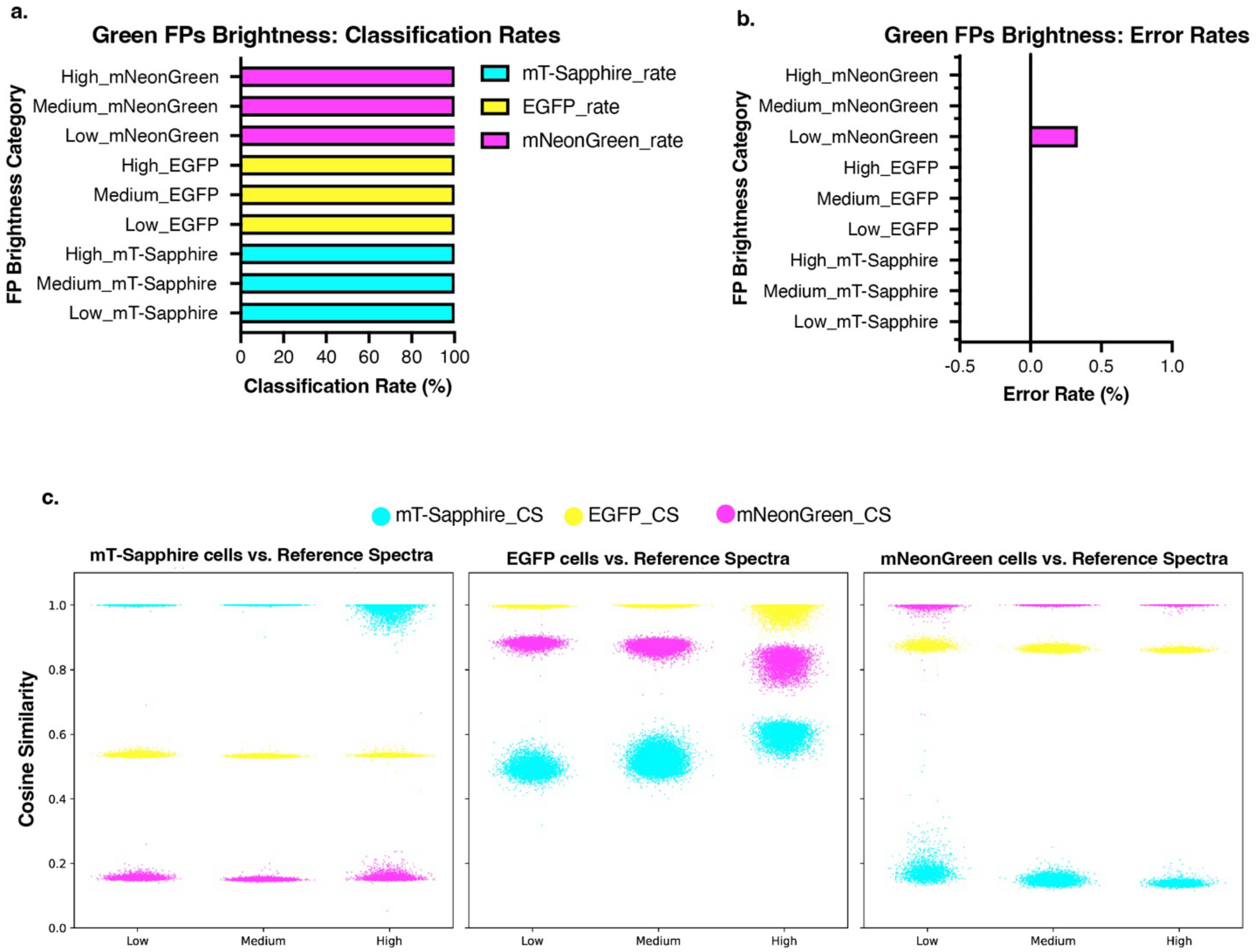


**Supplementary Figure 4. FP identity prediction is unaffected by signal intensity**. Individual cells within each reference spectra monoculture for mT-Sapphire, EGFP, and mNeonGreen were grouped into three brightness categories based on radiance quartiles (low, medium, high). **a-b.** The FP identity of each cell is predicted using MCS against all 3 green FPs reference spectra. The classification rate and error rates for each category are plotted. All cells are classified with 100% accuracy irrespective of brightness, with the exclusion of the mNeonGreen low brightness group (error rate of 0.33%). **c.** Plots showing cosine similarity values between cells and each FP reference spectra. Each FPs plot shows the data for cells based on intensity group.

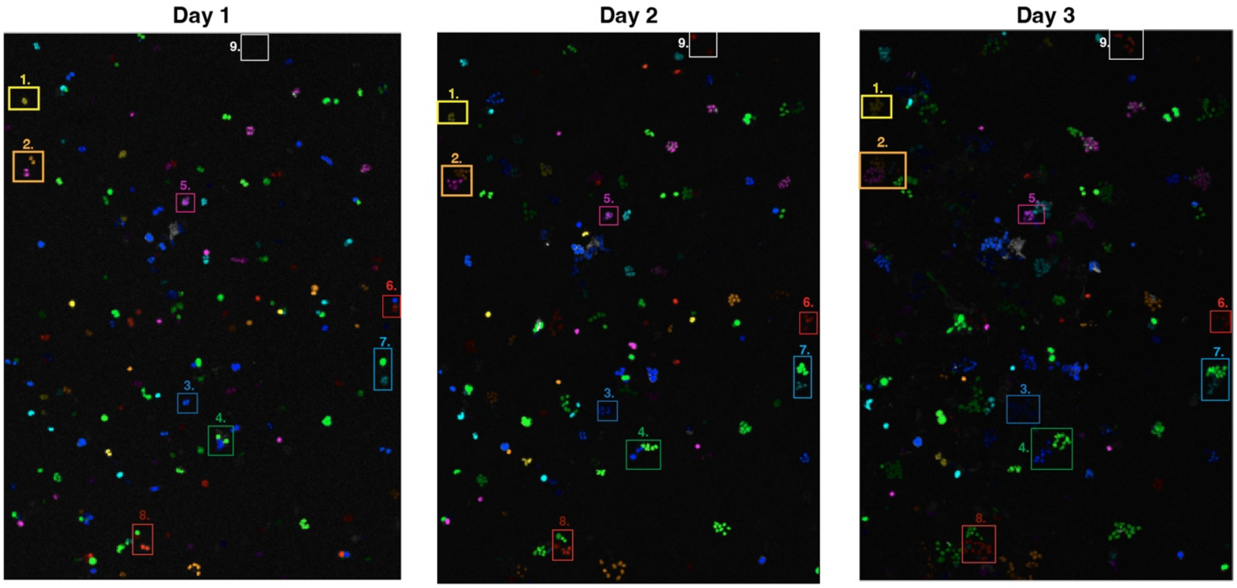


**Supplementary Figure 5. 3-day 12-Plex Clonal Growth.** Images taken on each 3 day of the 12-Plex clonal growth experiment. *CoSiDeX Paint-by-Numbers* tool used to pseudocolor cell masks with predicted FP identity. Rectangles show the representative clones shown in Figure 4h.

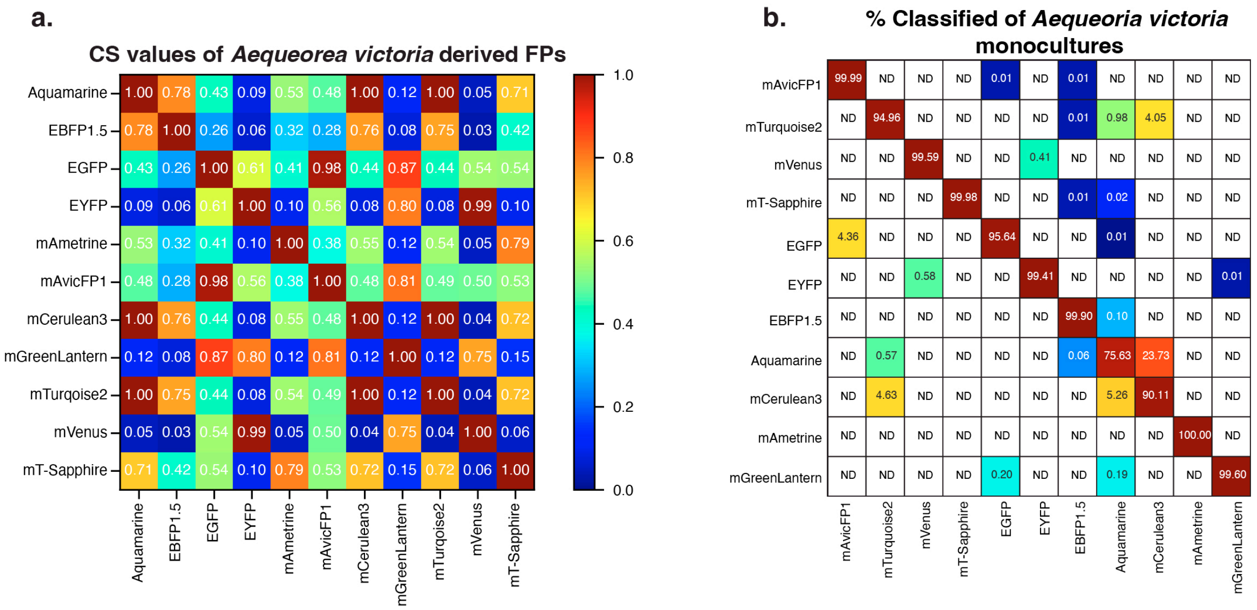


**Supplementary Figure 6. Resolvability of *Aequorea victoria* derived FPs. a.** Cosine similarity (CS) values of the 11 FPs originating from *Aequorea victoria*. A 0.9 CS threshold suggests that only a maximum palette of 7/11 FPs will be resolvable. **b.** Matrix showing the % of cells classified as each FP from the *Aequorea victoria* derived FP reference monocultures.

**Supplementary Tables 1-3 organized into a single .xlsx file.**

**Supplementary Table 1. Fluorescent protein properties extracted from FPbase.**

**Supplementary Table 2. Hyperspectral 4-Block imaging parameters**

**Supplementary Table 3. Hyperspectral reference spectra for 44 FPs**
